# Resolving the mesoscopic missing link: Biophysical modeling of EEG from cortical columns in primates

**DOI:** 10.1101/2022.03.16.484595

**Authors:** Beatriz Herrera, Jacob A. Westerberg, Michelle S. Schall, Alexander Maier, Geoffrey F. Woodman, Jeffrey D. Schall, Jorge J. Riera

**Author notes:** Denotes the corresponding author* **Corresponding Author** Jacob A. Westerberg, Department of Psychology, Vanderbilt University, 111 21^st^ Avenue South, 301 Wilson Hall, Nashville, TN 37240, USA. Denotes equal contribution (Beatriz Herrera, Jacob A. Westerberg).

## Abstract

Event-related potentials (ERP) are among the most widely measured indices for studying human cognition. While their timing and magnitude provide valuable insights, their usefulness is limited by our understanding of their neural generators at the circuit level. Inverse source localization offers insights into such generators, but their solutions are not unique. To address this problem, scientists have assumed the source space generating such signals comprises a set of discrete equivalent current dipoles, representing the activity of small cortical regions. Based on this notion, theoretical studies have employed forward modeling of scalp potentials to understand how changes in circuit-level dynamics translate into macroscopic ERPs. However, experimental validation is lacking because it requires *in vivo* measurements of intracranial brain sources. Laminar local field potentials (LFP) offer a mechanism for estimating intracranial current sources. Yet, a theoretical link between LFPs and intracranial brain sources is missing. Here, we present a forward modeling approach for estimating mesoscopic intracranial brain sources from LFPs and predict their contribution to macroscopic ERPs. We evaluate the accuracy of this LFP-based representation of brain sources utilizing synthetic laminar neurophysiological measurements and then demonstrate the power of the approach *in vivo* to clarify the source of a representative cognitive ERP component. To that end, LFP was measured across the cortical layers of visual area V4 in macaque monkeys performing an attention demanding task. We show that area V4 generates dipoles through layer-specific transsynaptic currents that biophysically recapitulate the ERP component through the detailed forward modeling. The constraints imposed on EEG production by this method also revealed an important dissociation between computational and biophysical contributors. As such, this approach represents an important bridge from the mesoscopic activity of cortical columns to the patterns of EEG we measure at the scalp.

**Highlights:**

- Cognitive EEG production was accurately modeled from empirically measured cortical activity in awake macaques.
- V4 laminar activity plausibly generates the attention-related signal indexed by the EEG.
- Models demonstrate the importance of biophysical geometry in cognitive EEG production.

## 1. Introduction

Identifying the neural sources of EEG is a substantial challenge facing the human neuroscience community. Many basic and clinical research programs rely on EEG as a window into human sensation, perception, cognition, and action. However, some insights into the mechanisms of these neural operations are invisible through this method. While the macroscopic EEG contains information, some of it is lost relative to the microscopic activations it reflects. As such, researchers are developing methods to extract as much information regarding the neural sources of EEG to strengthen this staple of neuroscientific approaches. EEG inverse source localization methods have offered a tool to estimate plausible brain sources generating such signals and, hence, make inferences about the underlying cortical mechanisms (Grech et al., 2008; Michel et al., 2004). Unfortunately, the results are indefinite because a given cranial polarization pattern can arise from multiple current configurations, requiring additional assumptions (Grech et al., 2008; Nunez et al., 2019). Researchers constrain the number of possible brain source configurations by imposing biophysical models to address this problem. The most widely used models are based on the equivalent current dipole, with each dipole representing the activity of a small cortical region (Nunez et al., 2019). The equivalent dipole model has been implemented in different inverse solutions, from a few discrete dipoles (Scherg et al., 2019) to more distributed inverse solution algorithms (e.g., sLORETA, MNE, MxNE) (Gramfort et al., 2012; Grech et al., 2008; Nunez et al., 2019).

Our goal is to supplement the anatomical constraints on EEG inverse solutions with biophysical and physiological constraints. This bolsters EEG’s interpretability regarding neural operations. We take a definite forward modeling approach. Detailed biophysical models of populations of neurons offer insights into the relationship between microscopic transmembrane currents and macroscopic cranial voltages through forward modeling (Cohen, 2017; Einevoll et al., 2019; Næss et al., 2021; Pesaran et al., 2018). Recent studies have linked transmembrane currents of pyramidal cells (PC) to equivalent dipoles and the dipoles to macroscopic EEG signals (Jones et al., 2009, 2007; Kohl et al., 2022; Law et al., 2022; Næss et al., 2021). Information derived from local field potentials (LFP) sampled across all the layers of the cerebral cortex can validate these equivalent dipoles (Murakami and Okada, 2006; Riera et al., 2012). A previous study in rat somatosensory cortex measured the current dipole moments from the laminar current source density (CSD) – a derivative of LFP – and described their contribution to concurrently recorded EEG (Riera et al., 2012). Although the strength of the current generator per surface area of active cortex arises from the current dipole moment density, it is invariant across brain structures and species due to common electrical biophysical properties of neurons (Murakami and Okada, 2015). As such, these considerations can generalize to model organisms more closely matching human cognition, such as the macaque monkey. Laminar recordings with CSD (Nicholson and Freeman, 1975; Pettersen et al., 2006) have already been used to elucidate the organization of multiple cortical areas in macaques (Bastos et al., 2020; Buffalo et al., 2011; Engel et al., 2016; Ferro et al., 2021; Godlove et al., 2014; Hansen and Dragoi, 2011; Hembrook-Short et al., 2017; Kajikawa et al., 2017; Klein et al., 2016; Maier et al., 2011, 2010; Mehta et al., 2000; Nandy et al., 2017; Ninomiya et al., 2015; Schroeder et al., 1998; Self et al., 2013; Tovar et al., 2020; Trautmann et al., 2019; van Kerkoerle et al., 2017; Westerberg et al., 2021, 2019) and their respective contributions to various EEG signals (Givre et al., 1994; Sajad et al., 2019; Westerberg et al., 2022).

Despite this progress, the biophysical relationship between the mesoscopic laminar LFP/CSD and the microscopic cellular sources has not been evaluated. This gap of knowledge renders the validity of predicting spatial patterns of cranial EEG voltage from current sources derived from laminar LFP questionable. Hence, we describe a biophysically plausible forward modeling approach linking mesoscopic CSD derived from LFP to macroscopic ERPs.

Based on earlier demonstrations that macaque monkeys produce homologues of human cognitive ERP components (Cohen et al., 2009; Heitz et al., 2010; Purcell et al., 2013; Reinhart et al., 2012; Sajad et al., 2019; Westerberg et al., 2020b; Woodman et al., 2007), it is now possible to obtain data necessary to measure current dipoles in the cerebral cortex under conditions generating cognitive ERPs. Recently, Westerberg et al. (Westerberg et al., 2022) reported current source density maps of the laminar distributions of transmembrane currents in extrastriate cortical area V4 associated with the cognitive ERP component known as the

N2pc. We use these data to demonstrate the utility of our detailed forward modeling approach. We show that V4 generates dipoles through layer-specific transsynaptic currents that biophysically contribute to ERP generation. Forward modeling this cortical activity renders EEG at the scalp consistent with that of previous reports of the representative ERP. Moreover, in evaluating the potential contributions of other cortical areas computationally involved in the ERP-indexed operation, this approach revealed that these computational contributors need not biophysically contribute to EEG production. In establishing a mesoscopic link between microscopic neural currents and macroscopic cranial voltages, this finding represents the first, definite forward model of a cognitive ERP from current dipoles derived from neural activity in primates.

## 2. Methods

### 2.1 Animal Care and Surgical Procedures

Procedures were in accordance with NIH Guidelines, AALAC Guide for the Care and Use of Laboratory Animals, and approved by the Vanderbilt IACUC following USDA and PHS policies. Two male macaque monkeys (*Macaca radiata*; monkey Ca, 7.5 kg; He, 7.3 kg) were implanted with MR compatible head posts and recording chambers with craniotomy over V4. One female macaque monkey (*Macaca radiata*; monkey Y, 7.3 kg) underwent an anesthetic event to perform anatomical imaging. Anesthetic induction was performed with ketamine (5–25 mg/kg). Monkeys were then catheterized and intubated. Surgeries were conducted aseptically with animals under isoflurane (1-5%) anesthesia. EKG, temperature, and respiration were monitored. Postoperative antibiotics and analgesics were administered. Further detail is documented elsewhere (Westerberg et al., 2020a, 2020b).

### 2.2 Magnetic Resonance Imaging

Anesthetized animals were placed in a 3 Tesla Magnetic Resonance Imaging (MRI) scanner (Phillips) at the Vanderbilt University Institute of Imaging Science. T1-weighted 3D MPRAGE scans were acquired with a 32-channel head coil equipped for SENSE imaging. Images were acquired using 0.5 mm isotropic voxel resolution with parameters: repetition 5 s, echo 2.5 ms, flip angle 7°.

### 2.3 Cognitive Task – Visual Search

Monkeys performed a color pop-out search. Search arrays were presented on a CRT monitor at 60 Hz, at 57 cm distance. Stimulus generation and timing were done with TEMPO (Reflective Computing). Event times were assessed with a photodiode on the CRT. We used isoluminant red and green disks on a gray background. Target feature varied within a session. Trials were initiated by fixating within 0.5 degrees of visual angle (dva) of a fixation dot. The time between fixation and array onset was between 750 and 1250 ms. A nonaging foreperiod function was used to determine the fixation period on a trial-by-trial basis. Arrays comprised of 6 items. Array item size scaled with eccentricity at 0.3 dva per 1 dva eccentricity so that they were smaller than the average V4 receptive field (RF) (Freeman and Simoncelli, 2011). The angular position of items relative to fixation varied session to session so that 1 item was positioned at the center of the RF. Items were equally spaced relative to each other and located at the same eccentricity. In each trial, one array item was different from the others. Monkeys made a saccade to the oddball within 1 second and maintained fixation within 2–5 dva of the target for 500 ms. Juice reward was administered following the successful completion of the trial. The target item had an equal probability of being located at any of the 6 locations. Eye movements were monitored at 1 kHz using a corneal reflection system (SR Research Eyelink). If the monkey failed to saccade to the target, they experienced a timeout (1–5 s).

### 2.4 Laminar CSD Recording Procedure

Laminar V4 neurophysiology was acquired at 24 kHz using a PZ5 and RZ2 (Tucker-Davis). Signals were filtered between 0.1-12 kHz. V4 data was collected from 2 monkeys (monkey Ca: left hemisphere; He: right) across 30 sessions (monkey Ca: 21; monkey He: 9) using 32-channel linear electrode arrays with 0.1 mm interelectrode spacing (Plexon) introduced through the intact dura mater each session. Arrays spanned layers of V4 with a subset of electrode contacts deliberately left outside of cortex. CSD was computed from the raw signal by taking the second spatial derivative along electrodes (Mehta et al., 2000; Nicholson and Freeman, 1975; Schroeder et al., 1998; Westerberg et al., 2019) and converting voltage to current (Logothetis et al., 2007). We computed the CSD by taking the second spatial derivative of the LFP:

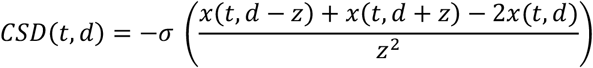

where *x* is the extracellular voltage at time *t* measured at an electrode contact at depth *d*, z is the inter-electrode distance, and *σ* is conductivity. CSD was baseline corrected at the trial level by subtracting the average activation during the 300 ms preceding array onset from the response at all timepoints. CSD was clipped 10 ms before saccade at the trial level to eliminate the influence of eye movements.

### 2.5 Laminar Alignment

Orthogonal array penetrations were confirmed online through a reverse-correlation RF mapping procedure (Cox et al., 2019; Nandy et al., 2017; Westerberg et al., 2019). RFs were found to represent portions of visual space consistent with previous reports of V4 (Gattass et al., 1988). An expanded description of the RF mapping procedure for this dataset has been reported previously (Westerberg et al., 2022, 2021). Positions of recording sites relative to V4 layers were determined using CSD (Nandy et al., 2017; Schroeder et al., 1998). Current sinks following visual stimulation first appear in the granular input layers of the cortex, then propagate to extragranular compartments. We computed CSD and identified the granular input sink session-wise. Sessions were aligned by this input sink. ‘L4’ refers to the granular input layer, ‘L2/3’ - supragranular layers, and ‘L5/6’ - infragranular layers.

### 2.6 Boundary Element Model

The monkey’s head was modeled as an isotropic and piecewise homogenous volume conductor comprised of the scalp, inner and outer skull, and the cortex surface. For the forward modeling of the experimental data, we employed the surfaces of the cortex, and the scalp and skull compartments obtained from the segmentation of the T1-weighted MRI of monkey Y in SPM12 (Penny et al., 2011) and BrainSuite (Shattuck et al., 2001), respectively. In the detailed biophysical simulations, we utilized the symmetric surfaces provided in the NMT v2 macaque atlas (Jung et al., 2021) to construct the head model (Figure 2A). The position of the EEG electrodes for both models was defined employing the EEG 10-10 system and the monkey’s scalp surface as described elsewhere (Giacometti et al., 2014). The scalp, skull, and brain conductivities were set as 0.43, 0.0063, and 0.33 S/m (Lee et al., 2015), respectively.

**Figure 1.**
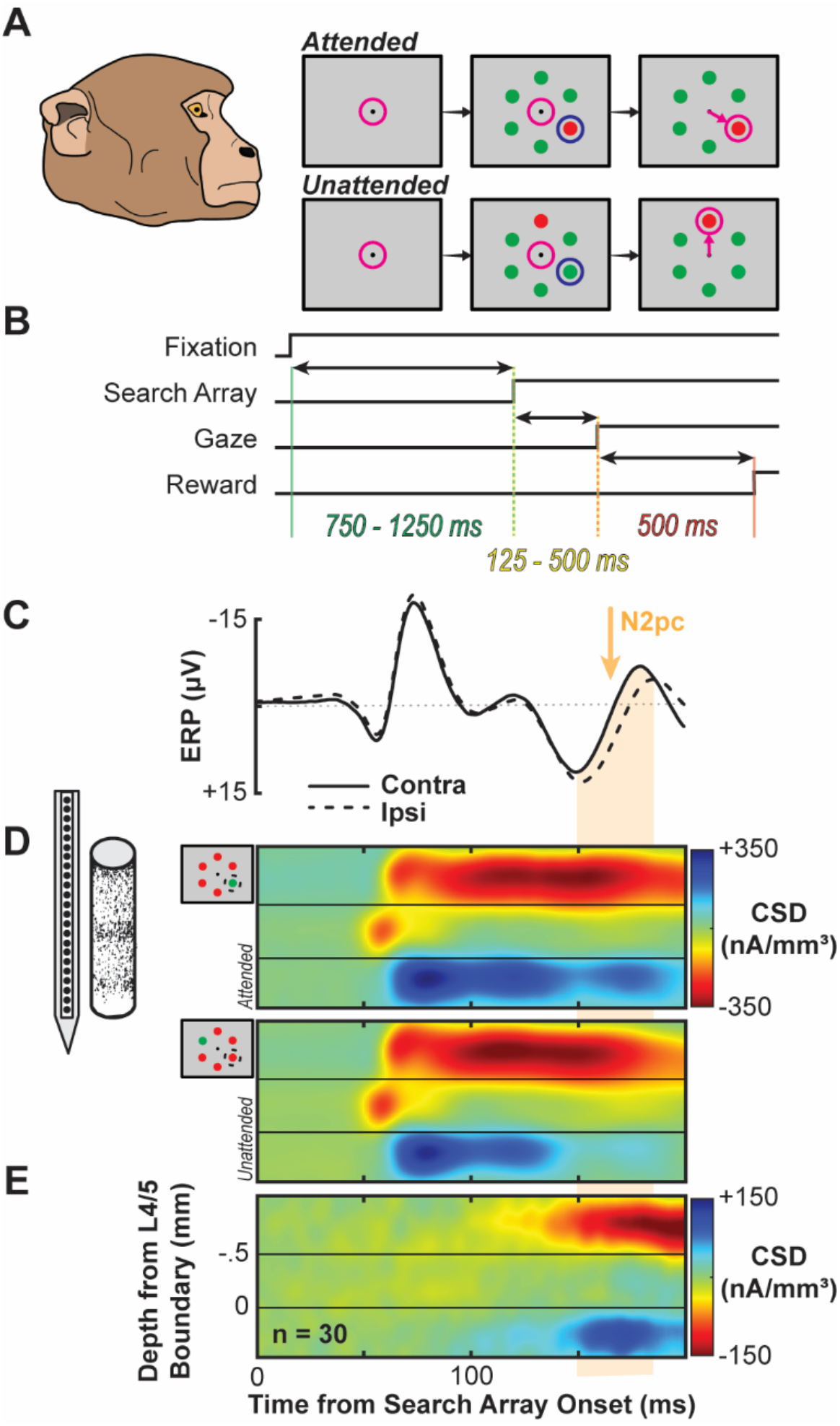
Experimental Design and Laminar Current Source Density. **A**. Monkeys viewed a fixation point and following a variable delay, a six-item visual search array appeared where one item was saliently different that the others (red among green or green among red). Monkeys shifted gaze to the oddball to receive a juice reward. Magenta circle indicates position of gaze. The oddball sometimes appeared in the receptive field (1/6 of trials) resulting in the attended condition. Blue circle indicates the position of the RF. All other trials were considered unattended. **B**. Timing of events in each trial. A dot appeared at the centered of the screen and once monkeys successfully made fixation, a 750-1250 ms delay ensued. The array appeared and monkeys made a saccade to the target as rapidly as possible (∼125 - 500 ms). Monkeys maintained fixation of the target for 500 ms to receive a juice reward. **C**. Proxy extracranial signal from an electrode placed outside the brain, above area V4. N2pc serves as our representative cognitive ERP indicating directed selective attention and can be observed ∼150-190 ms following search array onset and is highlighted in orange. **D**. Laminar current density computed across electrodes positioned along V4 layers averaged across sessions following alignment relative to the layer 4/5 boundary. Profiles shown for the attention condition (top) and unattended condition (bottom) and are plotted relative to the search array onset. **E**. Difference in current density profiles between attention conditions (attended – unattended) showing a difference in extragranular currents during the N2pc window.

**Figure 2.**
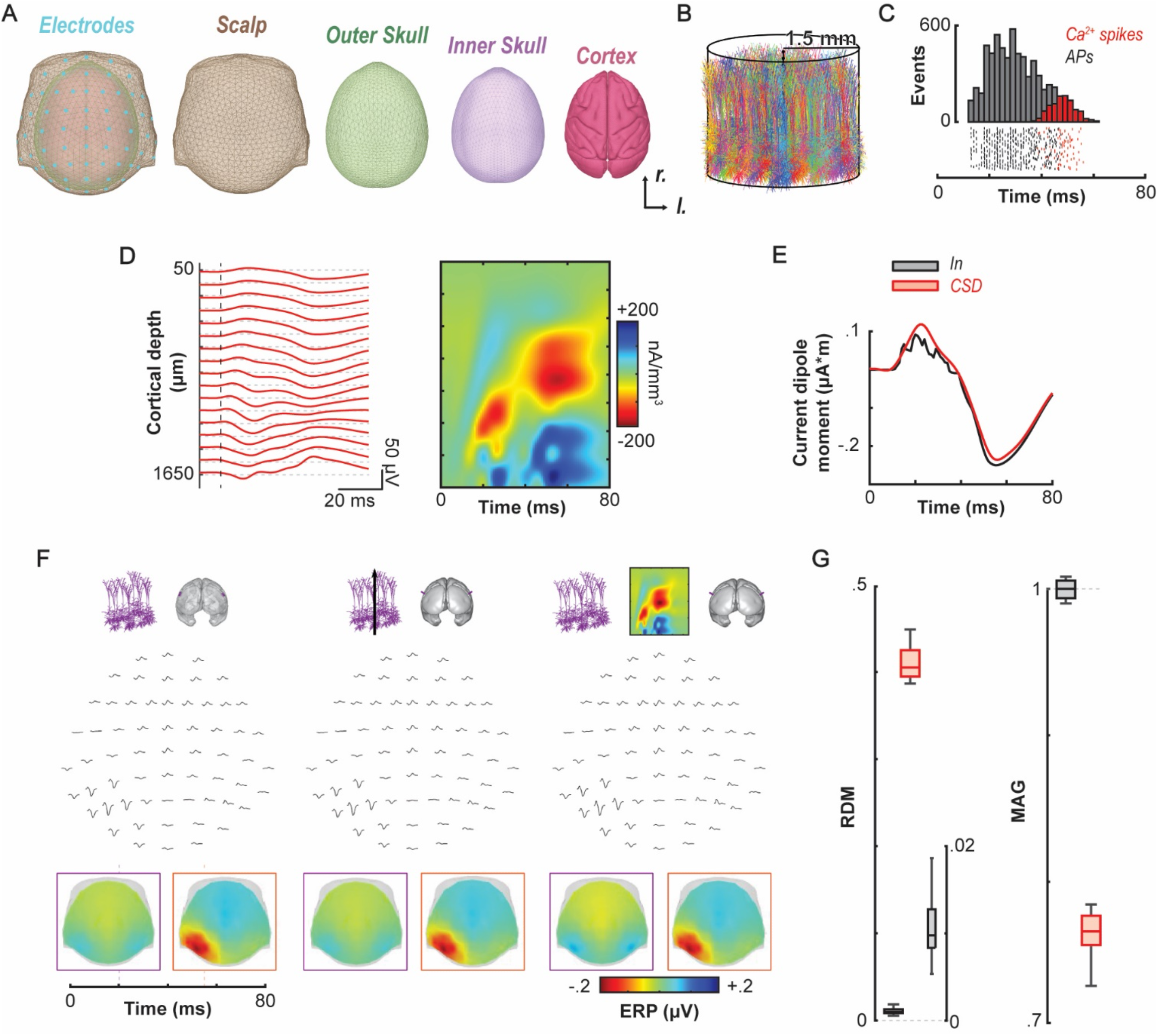
Biophysical forward modeling of synthetic data. **A**. Boundary element model composite and color-coded surfaces, obtained from the NMT v2 macaque atlas, used for the forward model. **B**. 2D representation of the simulated cortical column of 3 mm diameter formed by a collection of 1000 L5 pyramidal cells. The somas were randomly distributed in a 3 mm diameter cylinder and height corresponding to the approximate vertical extend of L5 in V4. **C**. Raster plots of 100 randomly selected neurons with the associated poststimulus time histogram for all neurons in the cortical column showing action potentials in black and dendritic Ca^2+^-spikes in red. **D**. LFPs and CSD evoked by suprathreshold stimulation of the collection of L5 pyramidal cells with a noisy current pulse (mean amplitude: 1.9nA). The stimulus onset for each neuron varied uniformly between 10ms and 20ms relative to the beginning of the simulation. **E**. Dipole moment calculated from the transmembrane currents of all neurons (In, black) and the estimated CSD (red). **F**. Simulated EEG resulting from the activity of two identical populations of 1000 L5 pyramidal cells under different stimulus located in V4, one in each hemisphere, obtained from each approach: detailed (left), exact dipolar (middle), and CSD dipolar (right), respectively. Neurons in both hemispheres were stimulated using the same stimulation paradigm as panels C-D. However, to simulate the asymmetry observed in the experimental recordings from V4, the mean amplitude considered for neurons in the right hemisphere was reduced to 1.85nA, decreasing the probability of eliciting dendritic Ca^2+^-spikes in these cells. Hence, reducing the amplitude of the late sink/sources present in the CSD map in panel D. The resulting spatial distribution of voltages at two time instances for each simulated EEG are shown at bottom. **G**. Relative difference – RDM – (left) and magnitude – MAG – (right) measures comparing the dipolar estimated EEGs to the ground-truth EEG (detailed approach) at 15 random locations in the monkey’s brain. RDM righthand ordinate: magnified view of the RDM for the exact dipolar approach (In).

### 2.7 EEG Forward Model

The EEG scalp potential 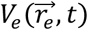 at any position 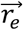 evoked by a continuous field of microscopic electric current sources 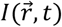 inside the brain *R* can be represented by an inhomogeneous Fredholm integral equation of the second kind (Riera et al., 2012):

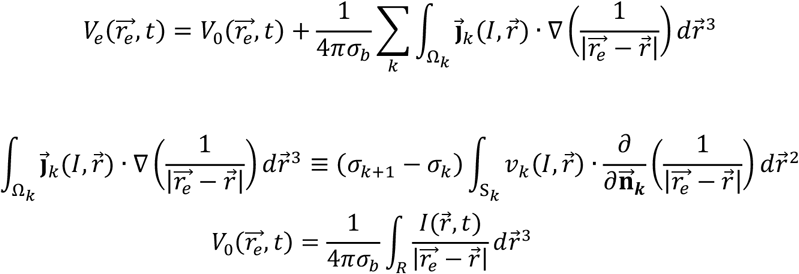

where *σ*_*k*_ denotes the conductivity of the *k-th* compartment in the head model (i.e., brain (*σ*_*b*_), skull, and scalp). 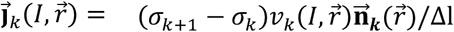 represents the secondary currents defined for each elemental volumetric shell Ω_*k*_ (i.e., a surface *S*_*k*_ of thickness Δ*l* → 0), and 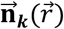 the normal vector to the surface *S*_*k*_ at location 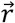.

We assume 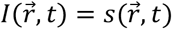 for 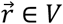 and 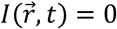 otherwise, where V is the volume of the brain region of interest, centered at 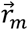 If the location of the EEG electrode 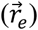 is far enough from the center 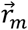, then 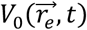 can be calculated as a function of the multipolar moments (Riera et al., 2012). Under the assumption of the dipolar model, the EEG forward model can be represented by the previous equation for 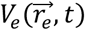 and the following equation for 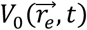:

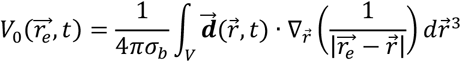

where 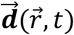 denotes the current dipole moment. The theoretical framework and numerical strategies used to compute the potentials 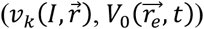 can be found elsewhere (Hamalainen and Sarvas, 1989).

The current dipole moment 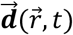 is assumed to originate from postsynaptic currents caused mainly by the activation of pyramidal cells perpendicular to the cortical surface (Hämäläinen et al., 1993) and from nonlinear processes taking place in their dendrites (Herrera et al., 2020; Suzuki and Larkum, 2017). Therefore, we can write 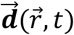 as 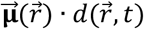, with 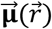 and 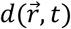 representing its orientation and time-varying amplitude, respectively (Riera et al., 2012). The activation waveform 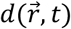 was estimated from the volumetric current sources (CSD) calculated from the laminar recordings, and the orientation of the dipoles from a weighted average of the normal to the cortical surface at the location of the dipole and the normal of its neighboring triangles. Considering the assumptions made to calculate the CSD, the amplitude 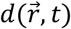 of the current dipole moment from the CSD in a volume of interest *V* is given by (Riera et al., 2012):

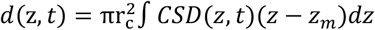

The z-axis is defined as the direction perpendicular to the cortex with positive and negative values toward the supragranular and infragranular layers, respectively. We modeled dipoles located in areas V4, lusV4, LIP, 7a, and FEF and assumed that the cortical columns in those areas were perfect cylinders of 3 mm radius (*r*_*c*_). The integrals were calculated using the trapezoidal method, where each subinterval corresponds to a particular grid point in the corresponding CSD method.

### 2.8 Biophysical Simulations

We generated a synthetic data set using detailed biophysical models of neurons to evaluate the equivalent current dipole derived from the CSD. We simulated the response to a noisy current pulse of 30 ms duration of a population of 1000 unconnected L5 pyramidal cells, described in a previously described model (Hay et al., 2011) (ModelDB, accession #139653, “cell #1”). The stimulus onset for each neuron was drawn from a uniform distribution, taking values between 10 and 20ms relative to the beginning of the simulation. The somas of the neurons were distributed uniformly in a cylinder of 3 mm diameter and height corresponding to the approximate vertical extend of L5 in V4, 1250, and 1750 *μm* below the pia matter (Figure 2B). All simulations were performed in Python using LFPy (Hagen et al., 2018), which builds on NEURON (Hines et al., 2009).

We calculated the LFP produced by the activity of the neurons at 17 equally spaced vertically aligned points located at the center of the cortical column, simulating a linear microelectrode array. As in the experiments, the inter-electrode distance was 100 *μm*. We employed the *point-source approximation*, implemented in LFPy, to compute the extracellular potentials with low-pass filtering at 100 Hz to obtain the LFP. The CSD patterns of the synthetic data sets were calculated using the spline-iCSD method (Pettersen et al., 2006) as implemented in the CSDplotter toolbox (https://github.com/espenhgn/CSDplotter) with custom MATLAB (R2021b, The MathWorks) scripts (Herrera et al., 2020).

To estimate the EEG produced by the simulated population of pyramidal neurons, we considered the current dipole moments calculated from the CSD, as explained in the previous section, and the transmembrane currents of all neurons in the cortical column. The equivalent current dipole produced by the simulated activity of the neurons can be estimated from the transmembrane currents of all neurons in the column as follows:

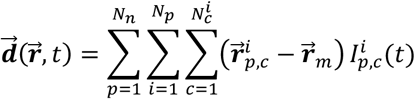

where *N*_*n*_, *N*_*p*_, and 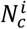 denote the total number of neuron types, the number of neurons in the *p-th* population and the number of compartments (*c*) in the *i-th* neuron of the *p-th* population, respectively. 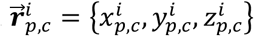 indicates the coordinates of the *c*-*th* compartment of the *i*-*th* neuron in the *p-th* population, 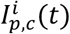 the transmembrane current of the *c*-*th* compartment of the *i*-*th* neuron in the *p-th* population and 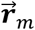 the center of mass of the cortical column, the coordinates of the dipole.

To compare the estimated EEG derived from these two approaches, we calculated the resulting EEG (ground-truth EEG) directly mapping the transmembrane currents into the scalp potentials described previously (Herrera et al., 2020). We quantified the mismatch error of each dipolar approach by calculating the magnitude (MAG) and the relative difference (RDM) measures (Meijs et al., 1989) between them and the ground-truth EEG at 15 randomly selected positions, designated as the center of the cortical column, throughout the monkey brain. A relative difference (RDM) closer to 0 and magnitude (MAG) closer to 1 indicates a higher degree of similarity between the dipolar approach and ground truth.

### 2.9 10-20 EEG Recordings

One monkey (monkey Z) was implanted with an array of electrodes approximating the human 10-20 system locations (FpFz, Fpz, F3, F4, Fz, Cz, C3, C4, Pz, P5, P6, POz, O1, O2, Oz) (Jasper, 1958). Referencing was done through linked ears. The impedance of the individual electrodes was confirmed to be between 2–5 kOhm at 30 Hz, resembling electrodes used for human EEG. EEG was recorded using a Multichannel Acquisition Processor (Plexon) at 1 kHz and filtered between 0.7–170 Hz. The monkey performed a visual search task necessitating directed spatial attention. Data was aligned to array onset and baseline corrected by subtracting the average activity during the 50 ms preceding the array onset from all timepoints. Data was clipped 20 ms prior to saccade to eliminate eye movement artifacts.

### 2.10 Comparing 10-20 EEG Recordings and Forward Models

To understand the configuration of plausible neural sources contributing to the generation of the N2pc, the N2pc measured in the 10-20 recordings was compared to different compositions of neural sources. 31 different configurations were generated with all possible combinations of V4, lusV4, LIP, 7a, and FEF sources. A single source model was simply the EEG modeled from a single source localized to that dipole location. Combination of source locations were computed as the sum of voltages from each source location for each simulated electrode site. In these models, we only considered the 15 electrode sites present in the 10-20 recording dataset rather than the full 10-10 configuration modeled elsewhere. Comparisons we performed as a Pearson correlation between each of the 15 empirical recording sites and the 15 simulated sites for each of the 31 model configurations. Therefore, each correlation comprised 15×30 data points. Whether a model was significantly correlated with the empirical measurement was evaluated with a Bonferroni corrected p value.

### 2.11 Data Availability Statement

The CSD data used to forward model EEG can be found through Data Dryad (https://doi.org/10.5061/dryad.djh9w0w15) or by request from the corresponding author. Code specific to the methods documented here can be obtained from the corresponding author.

## 3. Results

We introduce an approach for estimating the mesoscopic cortical columnar current dipoles from laminar *in vivo* field potential recordings to determine the contribution of distinct areas to a particular macroscopic EEG signal through forward modeling. The approach consists of the following steps: First, the current sources across depth were determined using CSD methods from laminar field potential recordings. Second, current dipole moments were calculated from the estimated CSD values. Third, the estimated current dipoles were used as current sources of forward models, accounting for their location and orientation. We validated this approach by simulating synthetic field potential recordings generated from detailed biophysical models of neuronal activity. This methodological framework was then applied to elucidate the neuronal generators of the ERP component known as the N2pc.

### 3.1 Intracranial current sources can be accurately estimated from LFPs

To evaluate the validity of intracranial current sources derived from CSD to identify cortical areas contributing to a spatial EEG pattern, we applied the new forward modeling approach to synthetic local field potentials generated using biophysical models of neuronal activity. Specifically, we simulated the activity evoked by a noisy current pulse in 1000 unconnected L5 pyramidal cells randomly distributed within a 3 mm diameter cylindrical cortical column (Figure 2B). Figure 2C-D shows the raster plots of 100 randomly selected neurons with the associated poststimulus time histogram of all simulated neurons and the associated laminar LFP and CSD. The resulting current dipole moments estimated from the transmembrane currents and the CSD are displayed in Figure 2E. Relative difference (RDM = 0.1447) and magnitude (MAG = 0.9571) measures suggest the two outcomes are in good agreement. Figure 2F illustrates the EEG estimated from each approach, generated from the activity of two identical columns in each hemisphere placed in a part of area V4 located on the lunate gyrus surface. Neurons in the two columns were stimulated with a noisy current pulse of different mean amplitude to mimic the asymmetry observed in the experimental recordings (Figure 3). The spatial distribution of voltages at two time points for each approach is shown at the bottom of Figure 2F. We compared the EEG derived from the two methods with the ground-truth EEG at 15 randomly selected locations throughout the monkey’s brain. We found that relative to the ground-truth EEG, the EEG derived from the pooled PC simulation corresponded better (mean ± SD RDM 0.0106 ± 0.0036; MAG 0.9996 ± 0.0062) than that derived from the CSD (RDM 0.4105 ± 0.0205; MAG 0.7634 ± 0.0183) (Figure 2G).

**Figure 3.**
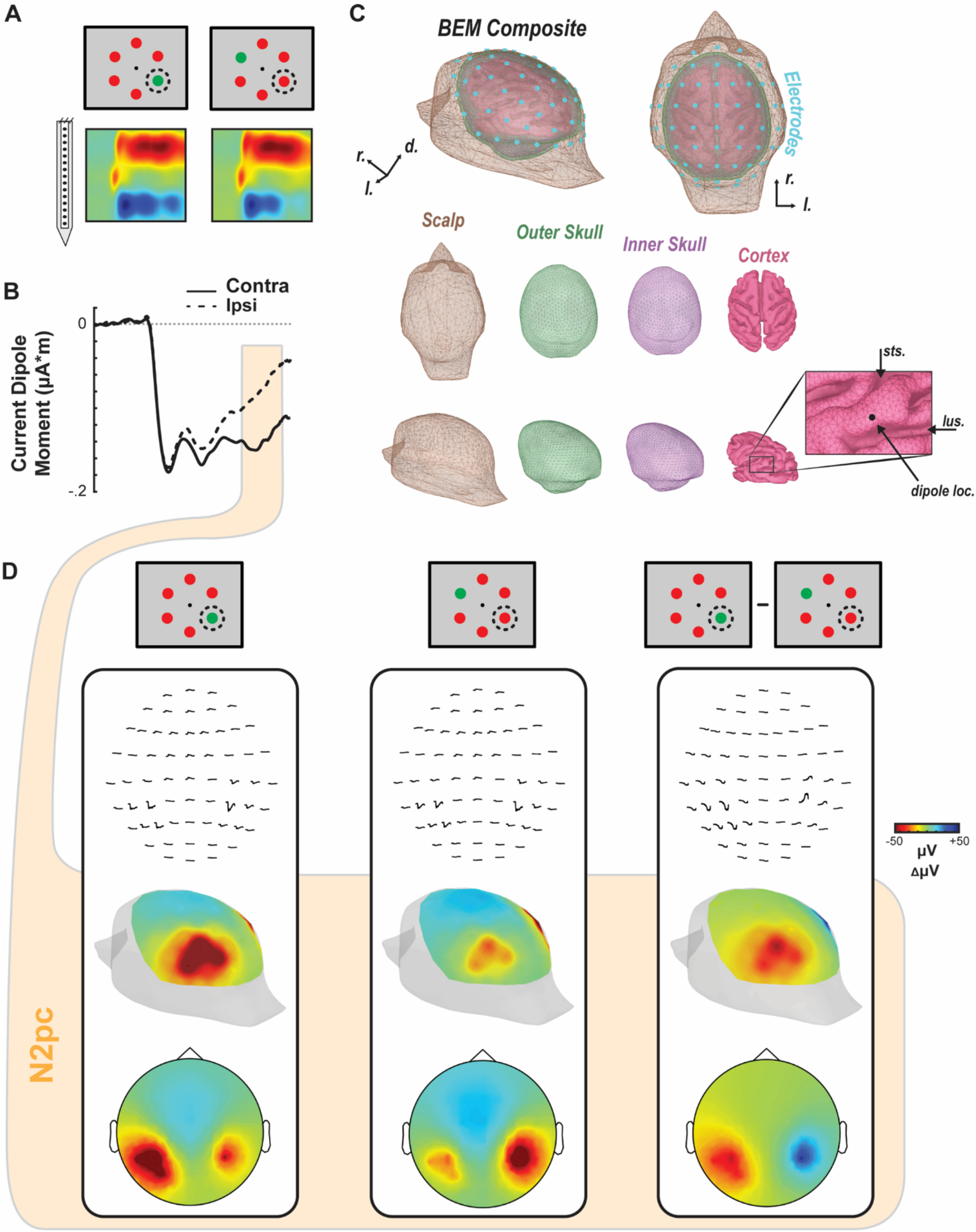
Biophysical forward modeling. **A**. Contrasting conditions (i.e., attended target in RF vs. attended target opposite RF) and corresponding CSD (n = 30 sessions) used for determining the current dipole moment used for the forward model. **B**. Dipole moment computed from the session averaged CSD (n = 30) across the time of array presentation for the target in RF (solid line, contra) and target opposite RF (dashed line, ipsi) conditions. N2pc epoch, defined as 150-190 ms after array onset, highlighted in orange. **C**. Boundary element model composite and color-coded surfaces, generated from MR scans from monkey Y, used by the forward model. Bottom right inset shows the position of the dipole used to generate the simulated EEG. Cyan discs (n = 61) represent the locations used to measure the simulated EEG. **D**. Simulated EEG distributions for a target inside the RF (left), target opposite RF (middle), and the difference between conditions (right). Voltage line plots across time for each simulated EEG electrode are shown at top. Voltage heatmaps for each condition are shown from a perspective over visual cortex (posterolateral view, center) and in 2D disc view (bottom). Heatmaps are plotted as average polarization during the time of N2pc (150-190 ms following array onset).

### 3.2 Experimental design

Before forward modeling, three objectives must be accomplished. First, monkeys must be trained to perform the cognitive task to proficiency. The representative ERP component used do demonstrate the novel forward modeling approach was the N2pc. The N2pc manifests as a function of the deployment of selective visual attention (Eimer, 1996; Luck and Hillyard, 1994, 1990; Woodman and Luck, 1999). To investigate the N2pc, we trained macaque monkeys to perform a visual search task (Figure 1A-B) requiring selective attention’s rapid deployment. Previous work demonstrates this task elicits an N2pc in macaque monkeys (Purcell et al., 2013). Two macaque monkeys (designated Ca, He) performed visual search for an oddball color target (red or green), presented within an array of 5 uniform distractors (green or red) (N sessions: monkeys Ca, 21; He, 9). Each animal performed above chance [chance level = 0.166] (behavioral accuracy: monkeys Ca, M = 0.88, SEM = 0.021; He, M = 0.81, SEM = 0.024) indicating they understood the task and were selectively deploying attention to accomplish it. Second, the ERP must be obtained to confirm it is being elicited. We sampled extracranial voltage fluctuations from an electrode placed outside the brain and above area V4. An N2pc was observed (Figure 1B) with this electrode. Third, suitable and replicable intracranial recordings must be obtained to measure local field potentials and calculate the current dipole during task performance. We obtained neural samples across the cortical layers with linear electrode arrays. After establishing that each penetration was orthogonal to the cortical surface and restricted to a cortical column, the synaptic activation was measured during task performance. The spatiotemporal profiles of synaptic currents are displayed in Figure 1C for both the attended condition (when the search target was present in the column receptive field) and the unattended condition (when the target was positioned outside the column receptive field). Recordings were restricted to area V4 on the prelunate gyrus, a hypothesized contributor to the N2pc (Westerberg et al., 2022; Woodman et al., 2007) and location where laminar activity orthogonal to the cortical surface can be reliably measured (Nandy et al., 2017). Coincident with the N2pc, we observed differences in synaptic currents across the layers of V4 (Figure 1D).

### 3.3 Modeling application to in vivo cortical activity

We employed forward modeling to compute the voltage distribution on the cranial surface caused by the translaminar currents in V4 (Nunez et al., 2019; Riera et al., 2012). Based on the average V4 current density (Figure 3A), we derived the dipole moment (Figure 3B) embedded in a boundary element model (BEM) of a macaque monkey head (Figure 3C). That dipole was strongest when the target was within the receptive field of the column. Lead fields were generated for a dipole on the convexity between the lunate and superior temporal sulci, where neurophysiological samples were taken (Figure 3C). Forward model solutions were calculated during the N2pc (150-190 ms following array presentation). The resulting spatial distribution of voltages was maximal posterolaterally with higher values contralateral to the target (Figure 3D). The spatial distribution of the difference in voltages when the target versus distractor was in the receptive field exhibited negativity contralateral to the target that was maximal posterolaterally. These results of the forward model were effectively indistinguishable from previous reports of the N2pc. Moreover, the forward model voltage distributions were robust to data thinning. That is, they were qualitatively similar when using the CSD from a single session (Figure 4A) as well as the CSD from each monkey individually (Figure 4B-C). Thus, currents measured in a column of V4 are sufficient to produce a voltage distribution over the scalp that emulates the observed N2pc.

**Figure 4.**
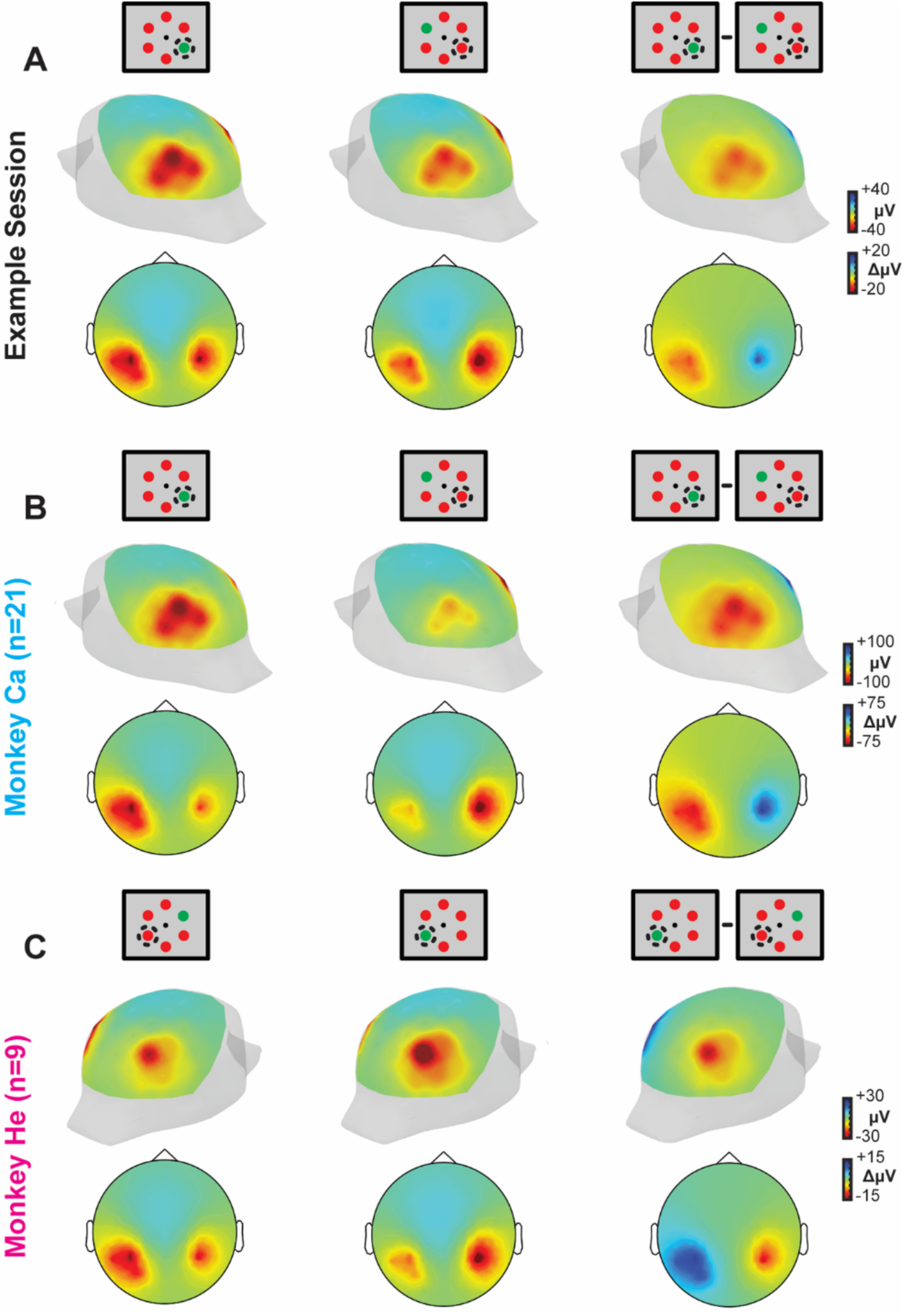
EEG forward models for example session CSD and individual monkey CSD. EEG voltage distribution over the scalp of the BEM simulated with a 61-channel electrode EEG array computed using CSD from an example session (**A**). and from CSD measured and averaged across sessions from monkey Ca (**B**) and monkey He (**C**). Both 3D posterolateral views and 2D disc views are shown for each subset of data. 3D head images are oriented such that the hemisphere in which the recordings were conducted are highlighted (left hemisphere, monkey Ca and example session; right hemisphere, monkey He). Note the difference plots are also computed such that the attention target ipsilateral to the recording chamber is subtracted from the contralateral presentations.

To investigate the specificity of this relationship, we calculated forward models produced by dipoles placed at other locations in V4. First, placing the dipole at two different locations on the convexity of the prelunate gyrus resulted in cranial voltage distributions that were qualitatively similar to one another as well as to the original dipole location (Figure 5A-B). This result confirms the robustness of the relationship between current dipoles in V4 and the N2pc. Second, the dipole was placed within V4 in the anterior bank of the lunate sulcus (Gattass et al., 1988) (Figure 5C). The dipole in this location is not oriented perpendicular to the cranial surface. Although LFP samples LFP have not been measured in this region, there is no *a priori* reason to assume that the laminar profile of sulcal CSD would be different from that on the gyrus within the same cortical region. The dipole in the sulcus, hereafter referred to as lusV4, resulted in spatial voltage distributions with more posterolateral distributions and similar lateralization relative to target hemifield. However, the forward model voltages were weaker than those derived from dipoles on the gyrus. These findings show that the summation of currents from multiple parts of V4, both gyral and sulcal, contribute to the N2pc. The spatially extensive appearance of the observed ERP thus can be partly understood as arising from variation in the orientation of contributing dipoles.

**Figure 5.**
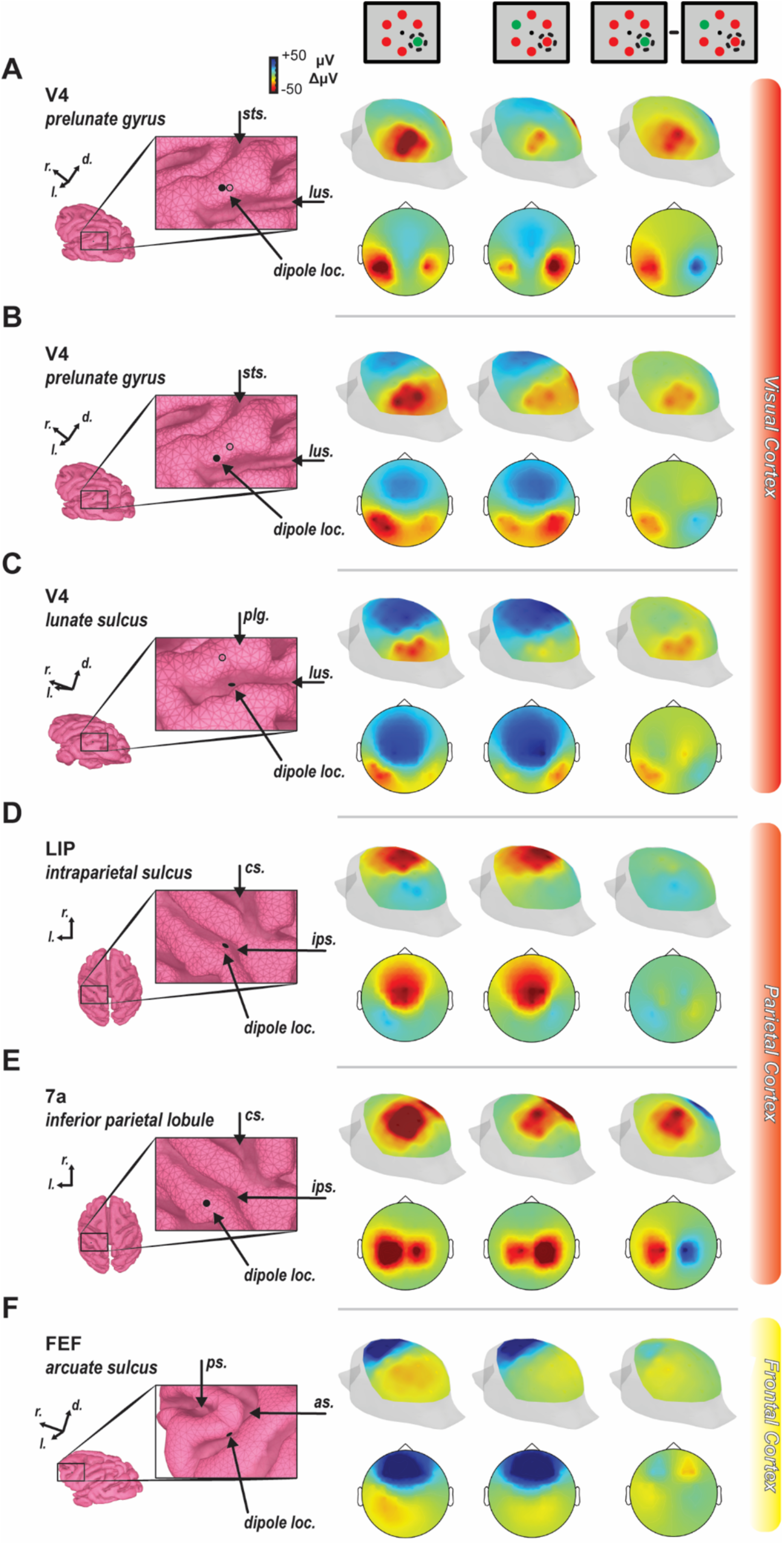
Forward models from different bilaterally symmetric dipole locations. The dipole was calculated from the current source density measured 150-190 ms following array presentation. Positions are shown on the enlarged cortex boundary element model surface with filled circle. In visual cortex panels, an additional unfilled circle denotes the position of the dipole used in Figure 2. Anatomical landmarks are indicated for reference in the inset images with the following abbreviations: sts – superior temporal sulcus, lus – lunate sulcus, plg – prelunate gyrus, cs – central sulcus, ips – intraparietal sulcus, ps – principal sulcus, as – arcuate sulcus. Cranial voltage heatmaps are displayed on posterolateral and 2D disc views for target in contralateral hemifield (left), target in ipsilateral hemifield (middle), and their difference (right). **A-B**. Dipoles placed at two other sites on the prelunate gyrus generate cranial voltages that were effectively indistinguishable from that observed with the first dipole location. **C**. Dipoles placed in part of V4 in the lunate sulcus generate a cranial voltage pattern with similar lateralization but different spatial distribution due to dipole orientation. **D**. The dipoles were also positioned in the lateral intraparietal area (LIP) on the lateral bank of the intraparietal sulcus, a previously hypothesized generator of the N2pc. **E**. Another location in parietal cortex (area 7a) was chosen as it also demonstrates robust attentional modulation and is located on a gyrus. **F**. Lastly, FEF was chosen as it has been related to the N2pc previously and also shows robust attentional modulation. It is important to note that the dipoles used throughout all panels were those measured in prelunate V4 and may or may not be a sufficient representation of the empirically measured currents in that area.

It seems unlikely that an ERP would arise from just one cortical area. Certainly, other cortical areas contribute to target selection and attention allocation during visual search. Using this forward modeling tool and assuming that the dipole measured in V4 is not too dissimilar from what would be measured in other cortical areas, we explored the possible contributions of other cortical areas to the N2pc. First, we modeled the possible contribution of two areas in parietal cortex. The lateral intraparietal area (LIP) is known to contribute to target selection during visual search and is located on the lateral bank of the intraparietal sulcus (Bisley et al., 2011; Goldberg et al., 2006; Ipata et al., 2006; Tanaka et al., 2015; Thomas and Paré, 2007). Area 7a, located on the inferior parietal lobule, also shows robust attention-related effects (Constantinidis and Steinmetz, 2001a, 2001b; Rawley and Constantinidis, 2010; Steinmetz et al., 1994; Steinmetz and Constantinidis, 1995), but unlike LIP, is biophysically oriented in a fashion more conducive to the generation of potentials measured at the scalp. These forward models assume that the laminar CSD in LIP and 7a is indistinguishable from that in V4, which is unknown but adopted provisionally. The dipole in LIP resulted in voltage with centromedial distributions and weak lateralization relative to target location. This confirms that neural processes in LIP contribute weakly if at all to the N2pc in macaques. Conversely, dipoles positioned in 7a produced robust potentials measurable at the scalp. However, these potentials were more centromedial than previous descriptions of the N2pc as well as the N2pc found in our own inverse modeling. These contrasting results demonstrate that some cortical areas can contribute to the microcircuit computational process indexed by a macroscopic ERP but need not contribute much biophysically to that ERP.

Third, we modeled the contribution of the frontal eye field (FEF) in the arcuate sulcus of the frontal lobe (Figure 5E). FEF plays a significant role in visual search and visual attention more generally (Schall, 2015), in part through influences on V4 (Moore and Armstrong, 2003; Moore and Zirnsak, 2017). Simultaneous recordings of spikes and LFP in FEF and occipital EEG have demonstrated correlations between LFP in FEF and N2pc polarization (Cohen et al., 2009; Purcell et al., 2013). The dipole in FEF resulted in cranial voltage distribution different from the observed N2pc but sharing a modest degree of lateralization relative to target hemifield. Thus, although FEF contributes computationally and neuroanatomically to the process indexed by the N2pc, it contributes little biophysically.

However, it is unlikely that a single area composes the entirety of any single EEG signal. Therefore, we modeled several combinations of plausible dipole sources and directly compared them to an empirically measured N2pc signal from a monkey with an array of 15 EEG electrodes positioned over the scalp (Figure 6A) performing the same visual search task across 18 recording sessions. To note, this third monkey (monkey Z) did not have any craniotomy which might impact the spatial spread of the EEG signal. Moreover, during the time period of the N2pc in our forward modeling data (150-190 ms after visual search array onset) we witness the same N2pc at posterior sites in the scalp EEG recording (Figure 6B). Significant difference was measured through a t-test on the difference between contra- and ipsilateral target presentations at sites P5 and P6 (t(35) = 2.42, p = 0.02).

**Figure 6.**
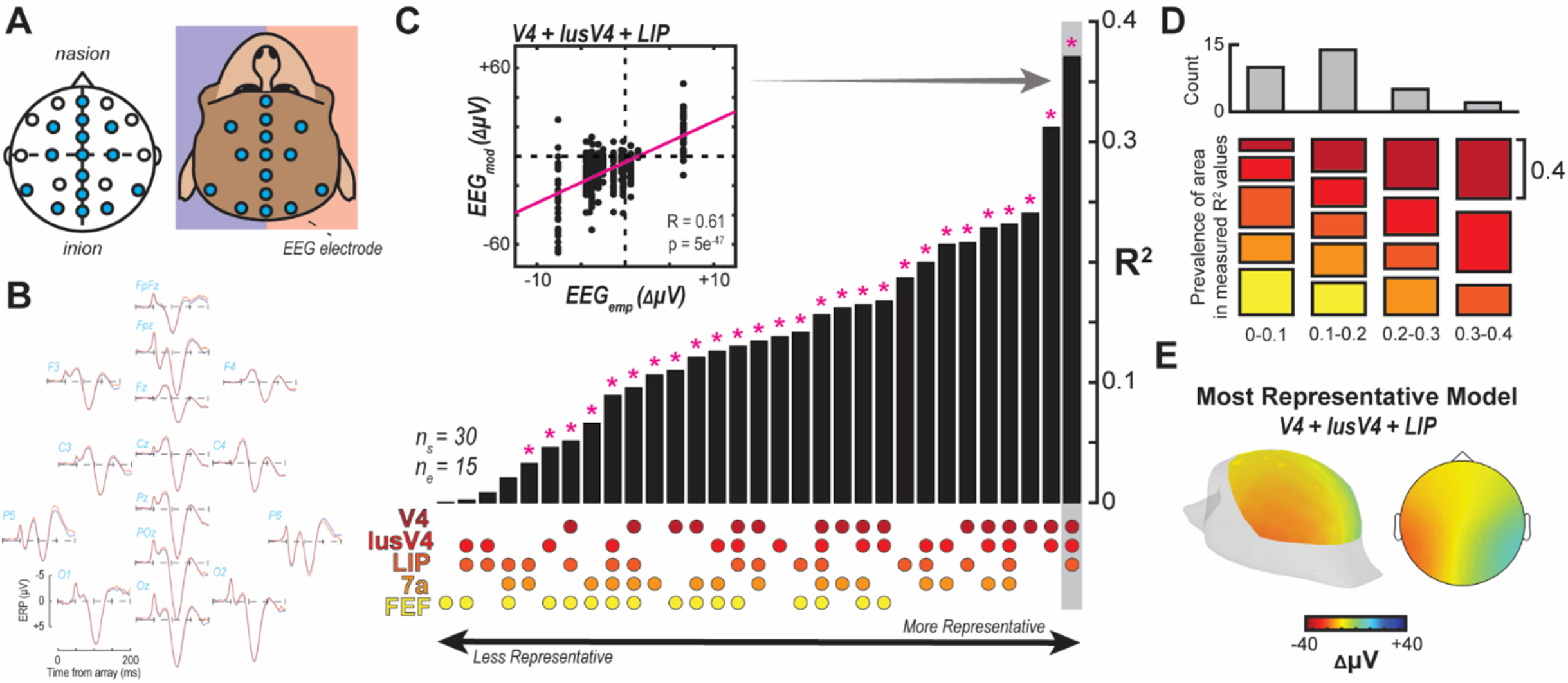
Comparison of N2pc voltage distribution over the scalp between empirical measurement and forward models. **A**. EEG was recorded in a subset of 10-10 configuration electrodes (n=15) from a monkey with intact skull. **B**. Voltage potentials for contra-vs. ipsilateral presentations of an attentional target in the visual search task. Differences are observed primarily at posterior sites about 150-190 ms following array presentation. **C**. Pearson correlation between EEG signal measured empirically (EEG_emp_) and the forward modeled EEG signal (EEG_mod_) for the difference between a contra-and ipsilateral target presentation at each of the electrode sites present in the empirical recording (n=15). Data was taken as the average difference between 150-190 ms following array presentation for both empirically measured and modeled data. EEG model was generated for each laminar recording session (n=30). Correlation was performed for each possible combination of V4, lusV4, LIP, 7a, and FEF sources (n=31). R-squared was computed and the bar plot reflects these values sorted from lowest to highest with significant correlation (following Bonferroni correction) indicated above with an asterisk. The combination of sources yielding each bar’s value is indicated below by dots with the legend at left. Inset shows the correlation observed with data for a single bar (V4+lusV4+LIP). **D**. R-squared values were grouped into 0.1 bins. Histogram plotted at top for the frequency of R-squared values. For each of those bins, the prevalence of each source was evaluated where the size of each colored bar indicates the total number of times that source was present in a model resulting in a R-squared value in the respective bin. For example, only V4, lusV4, and LIP sources were present in R-squared value measurements in the 0.3-0.4 bin. **E**. EEG distribution from the modeled data at the 15 recording sites using the most representative model (V4+lusV4+LIP).

For each of the 31 combinations of source locations, we measured the correlation between the empirical data and the modeled data at each of the matching 15 electrode positions. Note, a distinct forward model result was generated for each of the 30 laminar recording sessions meaning each correlation was between the empirical EEG voltage distribution and the 30 sessions of data for each of the 31 model configurations. A single example is shown in the inset of Figure 6C. Results of all correlations are summarized in Figure 6C with their corresponding configurations displayed below each bar. While many source configurations resulted in significant correlations with the empirical data, two stand out as superior to the others as indicated by their largest R-squared value – V4 + lusV4 and V4 + lusV4 + LIP. Note both of these models incorporate sources across both gyral and sulcal V4. Also, the better of the two also incorporates a parietal source. We found that the models have highest correlation with empirical values included gyral and sulcal V4 sources (Figure 6D). The most representative model exhibits the lateralized and posteriorly-distributed signature of the N2pc (Figure 6E).

## 4. Discussion

ERPs have been a powerful tool for investigating human cognition for decades. While insight can be gained and has been gained from these extracranial measures in isolation, some utility in these measures has yet to be extracted as uncertainty in their generation remains. In other words, knowing the composition of neural circuitry leading to the generation of specific cognitive signals in the EEG has the potential to inform what precisely the cognitive signal represents. However, measuring EEG generators directly in humans is unsystematic. Fortunately, cognitive EEG signals can be measured in macaque monkeys, a commonly used animal model for human cognition. Even so, a bridge from the neural activity of cortical columns, the purported generators of EEG signals, to the macroscopic EEG signal is missing. Here we provide an approach by which measurement of neural activity across the cortical layers in macaque monkeys during cognitive tasks is forward modeled to infer its impact on the global EEG signal. To our knowledge, this is the first documentation of forward modeling the synaptic currents across cortical layers to the EEG signal in awake, behaving primates.

In investigating the neural sources of the ERP component known as the N2pc as a demonstration for this approach, our results provide strong support for the hypothesis that it is generated from neural activity present in occipital and parietal areas. Early reports of the N2pc hypothesized that areas such as V4 contributed to its generation (Luck and Hillyard, 1994) with findings from an MEG study (Hopf et al., 2000) lending more direct support for occipital and parietal sources, specifically. Our comparison of various configurations of plausible neural sources with an empirically measured N2pc distribution suggests that the combination of sources in occipital area V4 and parietal area LIP best models the N2pc. This demonstration serves to highlight the importance of considering the biophysical geometry in EEG production. It is important to note that the dipole used for the modeling was measured in V4 and applied for all other areas in addition to V4. Therefore, neural recordings in LIP are necessary to determine whether a pattern of laminar activation sufficient to produce a dipole at the time of the N2pc is present. While that caveat for LIP remains, we can more confidently conclude that the laminar activation in V4 contributes to the N2pc as our measured activity comes specifically from that area.

While significant insight is gained from evaluating the biophysical plausibility of this variety of brain areas, it does highlight something rather unintuitive. That is, FEF and LIP – areas often synonymous with attentional control – on their own do not contribute much to the generation of the N2pc. In particular, a solo FEF source model is the least correlated with empirical data and FEF is not even present in the top 9-best correlated models. This highlights an important consideration in that the neural sources of computation (e.g., allocating attention or attentional modulation) and the neural sources of EEG generation need not be the same. That is, while an area like FEF has activation paralleling the N2pc (Cohen et al., 2009; Purcell et al., 2013), its physical position is not sufficient to contribute to its manifestation. In contrast, V4, an area showing more modest attentional modulation than FEF, is oriented in such a way that it is conducive to the generation of electric fields which can be measured at the scalp. However, it is entirely possible that the modulation of activity yielding the difference in dipole present in V4 producing the N2pc originates from FEF (Westerberg and Schall, 2021). Previous work has established that a robust set of connections exist between V4 and FEF (Anderson et al., 2011; Pouget et al., 2009; Schall et al., 1995; Stanton et al., 1995; Ungerleider et al., 2008) and modulation of FEF activity impacts V4 (Moore and Armstrong, 2003; Noudoost et al., 2014). Work remains to better understand the configuration of neural circuitry generating cognitive EEG signals and importantly, the origins of these signals – not just the areas producing the signal measured at the scalp.

While this study represents an important step in mapping microscopic through mesoscopic to macroscopic signals, this research program is by no means complete. Recent work highlights the strength of investigating the neural mechanisms of EEG at an even finer scale. We can better understand the generation of these potentials by understanding the underlying neural processes at the level of individual synapses through simulations (Herrera et al., 2020; Næss et al., 2021). In our modeling, we compute a dipole from the CSD, which is then used to calculate the definite voltage distribution at the scalp. This process sacrifices laminar information about the functional architecture generating EEG. For example, it is difficult to discern whether an observed source/sink in supragranular layers reflects synaptic activation onto the apical dendrites of L5 pyramidal cells, L2/3 pyramidal cells, or some combination (Pesaran et al., 2018). Additionally, CSD methods obscure any variation of neuronal activity in the radial plane of the column (Pettersen et al., 2006), parallel to the cortical layers.

Several decomposition techniques have been proposed to distinguish the contribution of distinct populations of neurons across layers to the LFPs (Di et al., 1990; Einevoll et al., 2007; Głąbska et al., 2014; Korovaichuk et al., 2010; Łęski et al., 2010; Makarov et al., 2010). Some methods are based on ICA and PCA decomposition methods that assume zero correlation or independence between the brain sources and do not account for the biophysical properties of area-specific cortical circuits (Di et al., 1990; Korovaichuk et al., 2010; Łęski et al., 2010; Makarov et al., 2010). Other techniques, such as the laminar population analysis (Einevoll et al., 2007) and dynamic causal modeling (Pinotsis et al., 2017), fit generative models, constructed from knowledge about local circuit processing and architecture to LFP recordings to estimate distinct intracranial sources. Yet, a systematic translation of these different neuronal sources to individual current dipoles and from them to macroscopic EEG signals is still lacking. Riera et al. (Riera et al., 2012) proposed a biophysical modeling strategy to link these scales. Their approach consisted of estimating the characteristic dynamic equations of intracranial currents from phenomenological models of principal neurons, pyramidal cells, and employing them to develop a generalized inverse solver that accounted for distinct neuronal population dynamics. The estimated brain sources would then be used to reconstruct the dynamics of principal neuronal populations through their phenomenological models. Alternatively, recent studies have inferred the circuit-level dynamics underlying cognitive ERPs by fitting the exogenous drives of a predefined canonical cortical microcircuit model to replicate the event-related current dipoles estimated through inverse modeling (Jones et al., 2009, 2007; Kohl et al., 2022; Law et al., 2022; Næss et al., 2021). The canonical cortical microcircuit model was constructed based on anatomical and electrophysiological findings from sensory cortical areas (Jones et al., 2009, 2007). New data from the agranular frontal cortex indicates that this microcircuit model does not generalize (Godlove et al., 2014; Ninomiya et al., 2015).

In sum, current research is rapidly generating more and more refined models of EEG production. Increasingly, the role of cortical columns – and the activity patterns they produce – is becoming clearer. However much of this is done computationally; experimental validation is necessary. Experimental elaboration of cortical columnar activity (e.g., relative contributions of cell types) seems the next step in following the computational work. However, the forward modeling we demonstrate here empirically resolves the missing link between theorized mesoscopic activity patterns to EEG production.

## Acknowledgments

This work was supported by NEI [grant numbers: P30EY008126, R01EY019882, R01EY008890, R01EY027402] and the Office of the Director [grant number: S10OD021771]. J. A. W. was supported by fellowships from NEI [grant numbers: F31EY031293, T32EY007135]. The authors would like to thank B. Williams, D. Richardson, I. Haniff, L. Toy, M. Feurtado, M. Maddox, R. Williams, and S. Motorny for technical support. The authors would like to thank A. Sajad, E. Sigworth, K. Lowe, S. Errington, T. Reppert for helpful conversations regarding the work.

## Author Contributions

B. H., J. A. W., M. S. S., A. M., G. F. W., J. R. R., and J. D. S designed the research. B. H. and J. A. W. analyzed the data. J. A. W. performed research. J. A. W. prepared visualizations. B. H., J. A. W., M. S. S., A. M., J. R. R., G. F. W., and J. D. S. wrote the manuscript.

## Declaration of Interests

The authors declare no competing interests.

## Notes

### Competing Interest Statement

The authors have declared no competing interest.

